# *De novo* designed pMHC binders facilitate T cell induced killing of cancer cells

**DOI:** 10.1101/2024.11.27.624796

**Authors:** Kristoffer Haurum Johansen, Darian Stephan Wolff, Beatrice Scapolo, Monica L. Fernández Quintero, Charlotte Risager Christensen, Johannes R. Loeffler, Esperanza Rivera-de-Torre, Max D. Overath, Kamilla Kjærgaard Munk, Oliver Morell, Marie Christine Viuff, Alberte T. Damm Englund, Mathilde Due, Stefano Forli, Emma Qingjie Andersen, Jordan Sylvester Fernandes, Suthimon Thumtecho, Andrew B. Ward, Maria Ormhøj, Sine Reker Hadrup, Timothy P. Jenkins

## Abstract

The recognition of intracellular antigens by CD8^+^ T cells through T-cell receptors (TCRs) is central to adaptive immunity, enabling responses against infections and cancer. The recent approval of TCR-gene-edited T cells for cancer therapy demonstrates the therapeutic advantage of using pMHC recognition to eliminate cancer. However, identification and selection of TCRs from patient material is complex and influenced by the TCR repertoire of the donors used. To overcome these limitations, we here present a rapid and robust de novo binder design platform leveraging state-of-the-art generative models, including RFdiffusion, ProteinMPNN, and AlphaFold2, to engineer minibinders (miBds) targeting the cancer-associated pMHC complex, NY-ESO-1^(157-165)^/HLA-A*02:01. By incorporating *in silico* cross-panning and molecular dynamics simulations, we enhanced specificity screening to minimise off-target interactions. We identified a miBd that exhibited high specificity for the NY-ESO-1-derived peptide SLLMWITQC in complex with HLA-A*02:01 and minimal cross-reactivity in mammalian display assays. We further demonstrate the therapeutic potential of this miBd by integrating it into a chimeric antigen receptor, as *de novo* Binders for Immune-mediated Killing Engagers (BIKEs). BIKE-transduced T cells selectively and effectively killed NY-ESO-1^+^ melanoma cells compared to non-transduced controls, demonstrating the promise of this approach in precision cancer immunotherapy. Our findings underscore the transformative potential of generative protein design for accelerating the discovery of high-specificity pMHC-targeting therapeutics. Beyond CAR-T applications, our workflow establishes a foundation for developing miBds as versatile tools, heralding a new era of precision immunotherapy.

## INTRODUCTION

T cells are immune cells capable of responding to intracellular antigens, for example, derived from infections or cancerous mutations. T-cell antigen recognition relies on a T-cell receptor (TCR) that recognises a peptide presented by a major histocompatibility complex (pMHC) on the surface of antigen-presenting cells. This recognition is primarily driven by Major Histocompatibility Complex class I (MHC-I) surface expression and CD8^+^ T cells expressing a TCR that can recognise the given pMHC complex; this identification of TCRs for therapeutic use is driven by selection of therapeutically relevant T-cell populations from patients^1,2^ or driving the expansion of a T-cell population from the naive repertoire towards a given antigen in healthy donors^3^. These approaches are both laborious and technically challenging, and potentially limited or biased by the TCR repertoire in the donors used for selection. Furthermore, it remains challenging to predict the behaviour and potential cross-reactivity of such TCRs when applied in hosts other than the donors they are selected from^4,5^.

To facilitate immunotherapies based on pMHC recognition, strong interest has emerged in developing alternative strategies to target intracellular proteins via the pMHC complex, avoiding the challenges associated with TCR identification. TCR-like molecules, which include TCR-like antibodies, TCR bispecific engagers and single-chain TCRs^6^, are promising approaches. Such TCR-like molecules can be incorporated into chimeric antigen receptors (CARs) to induce T cell-mediated cytotoxicity towards a given pMHC target^7^. However, TCR-like antibody generation relies either on mouse hybridoma technology or phage display screening. Whilst some success has been reported, these approaches often struggle to generate high-affinity binders. Hybridoma technology is limited by low throughput and labour-intensive protocols, and it skews towards MHC rather than peptide specificities^8,9^. Phage display panning is faster and more cost-effective than hybridoma technology, but the libraries still rely on large naïve antibody repertoires^10^. Alternative methods have been explored *in silico* and *in vitro*^11,12^ but are limited by previous knowledge of TCR binding interactions. Furthermore, establishing the specificity of high-affinity binders via exploration of cross-reactive binding presents a substantial challenge for current *in vitro* approaches^13^. Fortunately, recent developments in computational structural modelling pave the way for new opportunities for *in silico* exploration of the space of TCR-like pMHC binders. It was recently demonstrated that structural models such as TCRen^14^ or DeepAIR^15^ could predict TCR-pMHC interaction with remarkable accuracy. Despite this progress, structural analysis has been overshadowed by the unprecedented advances in generative protein design.

Up until a year ago, machine learning-guided protein design required the expensive screening of tens of thousands of variants for only a few binders. Benefiting from advances in image AI, the rise of denoising diffusion models is now beginning to unlock the full potential of generative binder design. Among various *de novo* protein design approaches, RFdiffusion is considered one of the most promising tools due to an impressive average of ∼10% success rate of designed binders presenting ultra-high target affinity (<10nM)^16^. The design pipeline consists of three independently trained models, namely the diffusion model, which *de novo* generates an highly accurate scaffold, followed by ProteinMPNN^17^, and finally, AlphaFold2 (AF2), which can predict and evaluate a protein structure by its sequence^18,19^. Here we show that these advances in *de novo* binder design, screening, and refinement of specific pMHC-binding minibinders (miBds) can be applied to rapidly generate specific binders to pMHCs. We demonstrate that these can be readily adapted for CAR constructs; a concept we call *de novo* pMHC-Binders for Immune-mediated Killing Engagers (BIKEs), allowing for the engineering of T cells to mediate killing of antigen-expressing cancer cells through pMHC recognition.

## MAIN

### Design and *in silico* cross-panning of NY-ESO-1 binders

MHC-I molecules, responsible for presenting intracellular peptides to cytotoxic CD8^+^ T cells, adopt a globular structure composed of three extracellular domains (α1, α2, α3) and a β2-microglobulin (β2m) subunit. The α1 and α2 domains form a peptide-binding groove, which accommodates short peptides (typically 8-11 amino acids) and displays them for recognition by TCRs. In our work, we focused on designing binders that specifically target the pMHC formed by MHC-I, presenting the cancer-associated NY-ESO-1 peptide SLLMWITQC (NY-ESO-1^(157-165)^) in the context of HLA-A*02:01 (SLLMWITQC/HLA-A*02:01)^20^. NY-ESO-1, also known as Cancer/Testis Antigen 1B (CTAG1B), is a well-described cancer-testis antigen which elicits T-cell recognition and is expressed in a wide range of tumours, including melanoma, lung, ovarian, and breast cancers, but not in normal tissues, except for the testis (Figure 1a). This broad, cancer-specific expression pattern makes it an attractive target for cancer immunotherapy. We chose to target our design efforts against non-anchor residues of the peptide to ensure binding focus to the peptide and not just the MHC class I. Following backbone generation through RFdiffusion denoising trajectories, we designed sequences using ProteinMPNN (Figure 1b). The resulting designs were filtered based on AF2 *initial guess* (ipAE cut-off of 12 Å and a pLDDT cut-off of 88), then partial diffusion and ProteinMPNN were used for those designs in an attempt to in silico affinity mature the designs. Designs that then passed an ipAE cut-off of 7 Å and a pLDDT cut-off of 92 were selected for *in silico* cross-panning. Here, we again leveraged AF2 *initial guess* to generate binding scores for the top binders not only to the target pMHC complex but also 10 mutants of the peptide in HLA-A*02:01 modelled with Alphafold3^21^ (Figure 1c). Nine variant peptides were close variants (single or double mutants) of the wild-type SLLMWITQC sequence, while one was a quadruple mutant. Briefly, the binders with low ipAE score (<10) for the target and high (>10) scores for the nine variant peptide/HLA-A*02:01 complexes were preferentially selected, with the highest predicted specificity observed for NY1-B04 (Figure 1c). We reasoned that this approach could increase the chance of binder specificity and reduce downstream off-target effects. We observed that all of the pMHC miBds were predominantly composed of multiple α helices with short linking regions classified as β sheets (Supplementary Figure S1, Extended Data Table 1). These helices were typically arranged into four tightly packaged units, aligning parallel or near-parallel to the flanking α helices of the MHC-I, encapsulating the peptide (Supplementary Figure S1c). Attempts to build a binary logistic regression classifier to predict an overall orientation of the miBds was hindered due to the limited dataset size and variation. This made robust training and testing challenging. The C- and N-termini were oriented parallel to the peptide and near each other but did not directly interact with the peptide. The 44 binders with the best scores across all targets were selected for experimental validation.

**Figure 1.**
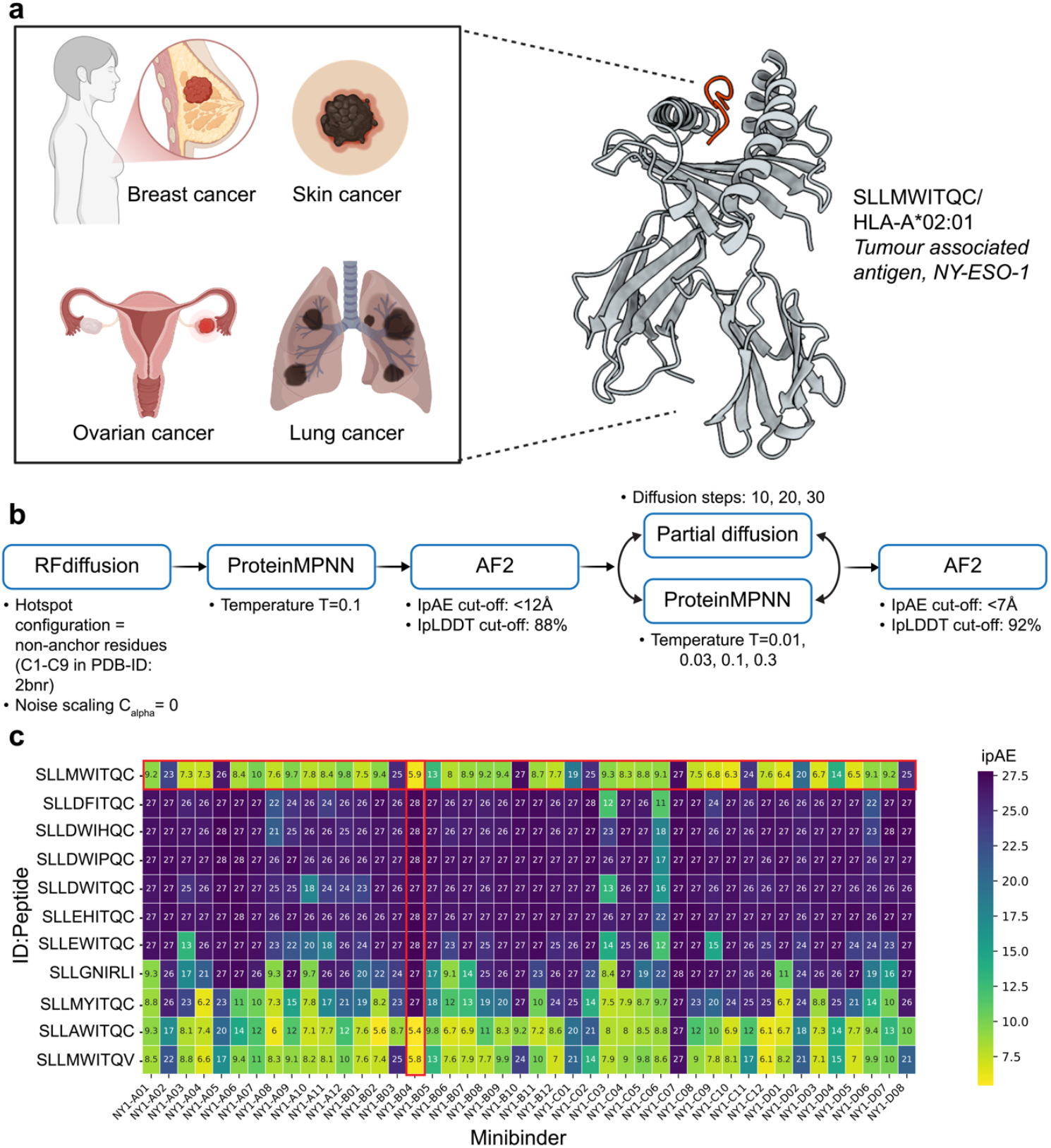
Design of minibinders (miBds) against NY-ESO-1^(157-165)^ presented on pMHC. **(a)** NY-ESO-1 (SLLMWITQC/HLA-A*02:01) is a highly immunogenic cancer antigen expressed in a wide range of tumours, including melanoma, lung, ovarian, and breast cancers, but only expressed in testis of healthy tissues. **(b)** Pipeline for identification of miBds for the SLLMWITQC/HLA-A*02:01 pMHC complex with specifications indicated. **(c)** Heatmap showing ipAE scores as parameter for potential binding interaction of the 44 miBds selected for *in vitro* characterisation towards the original SLLMWITQC peptide and 10 selected variant peptides. Red squares mark the target SLLMWITQC peptide (horizontal) and the NY-B04 miBd (vertical).

### Molecular dynamics (MD) identification of false positives and off-target binding likelihood

Although recent advances in the protein design and structure prediction field have revolutionised the design and prediction of *de novo* miBd complexes, there is still a high rate of false positives, where predicted binders fail during experimental testing. Thus, identifying descriptors that can facilitate the downselection of miBd-target interfaces to decrease the false positive rate is crucial. Protein-protein binding is determined by an interplay of enthalpy and entropy, and therefore, when characterising and predicting protein-protein interfaces, considering both components is critical. We found that conformational diversity is a key determinant to facilitate the selection of potential binders and to identify false positives (Figure 2a-c). To define the protein-protein interfaces for interface B-factor, electrostatic, and Van der Waals interaction energy fluctuations, we considered the salt-bridge and hydrogen bond interactions for the miBd-pMHC complexes that occur more than 10% throughout the simulations (Figure 2a). The AF2-predicted complex highlights critical salt bridge and charged hydrogen bond interactions of NY1-B04 with HLA-A*02:01 and shows the interactions of NY1-B04 with the SLLMWITQC NY-ESO-1 peptide (Figure 2b). In particular, residues Glu11, Glu96, Thr133, Arg139 and Glu140 of NY1-B04 are crucial interaction partners of HLA-A*02:01, and residues Tyr107, Glu115, Gln119, and Lys212 play a key role in stabilising the peptide. The radar plot based on these three descriptors shows that miBd NY1-B04 revealed the lowest conformational flexibility in the binding interface and the lowest interaction energy fluctuations, representing the most promising miBd-pMHC complex within the dataset (Figure 2c). As such, we hypothesised that it was the most likely to bind the target pMHC but proceeded to test all 44 miBds to explore this prediction.

**Figure 2.**
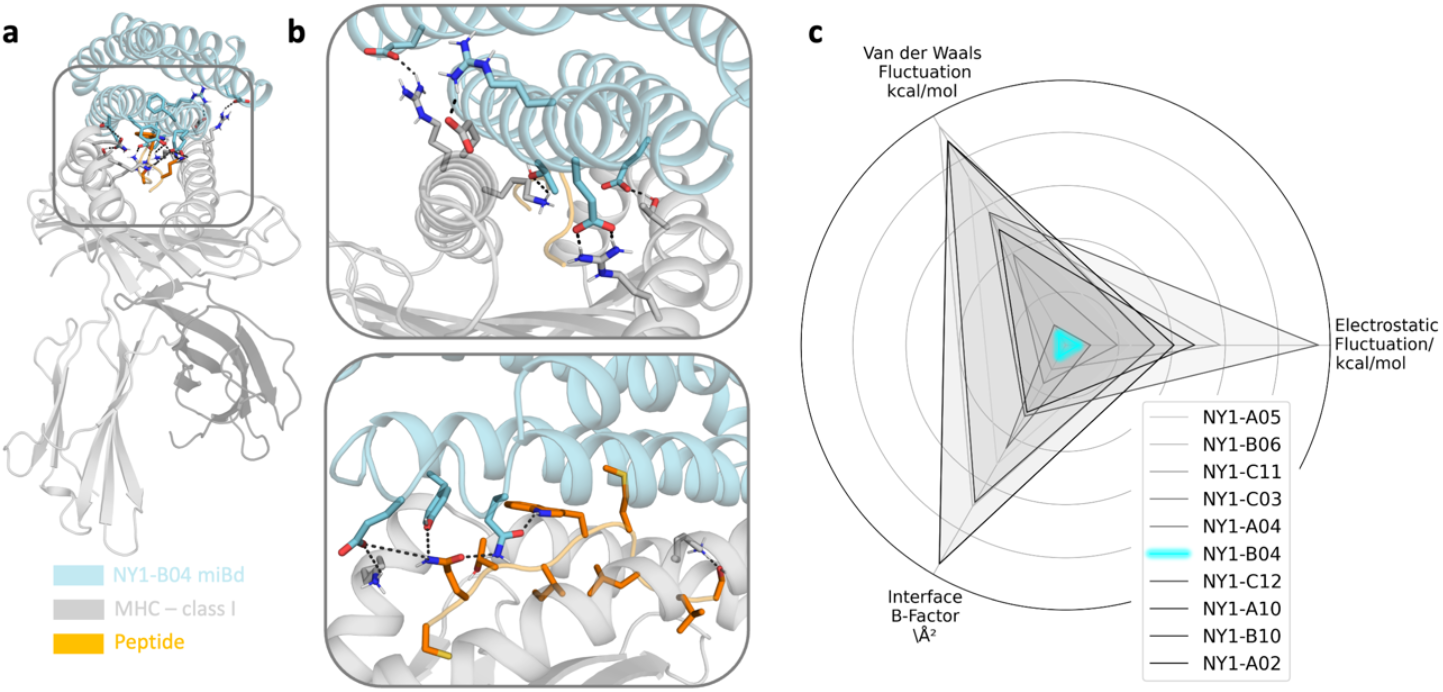
Conformational diversity estimators from MD simulations are able to downselect the number of false positives and predict pMHC binder. **(a)** Characterisation of dominant salt-bridge and hydrogen bond interactions involved in stabilising NY1-B04 with the target pMHC SLLMWITQC/HLA-A*02:01. **(b)** Close-up highlighting the salt bridge and charged hydrogen bond interactions of NY1-B04 (cyan) with HLA-A*02:01 (grey) (top) and the interactions of the peptide (orange) SLLMWITQC with NY1-B04 (bottom). **(c)** Fluctuations in electrostatic and van der Waals interactions combined with an interface-residue-B-factor analysis as estimators for conformational flexibility in the binding interface identify miBd NY1-B04 (cyan) as the most promising miBd-pMHC complex.

### *In vitro* selection and specificity screening of SLLMWITQC/HLA-A*02:01 binders

To experimentally validate binding and specificity towards pMHC, synthetic genes encoding all 44 miBd designs targeting SLLMWITQC/HLA-A*02:01 were cloned into a lentiviral CD28-CD3ζ CAR vector for expression on the cell surface (Figure 3a). Using a mammalian surface display (MSD) system based on miBd-CAR transduced Jurkat cells, we evaluated the binding capacity of the miBd library to SLLMWITQC/HLA-A*02:01 through pMHC-tetramer binding (Figure 3a-b). From this library, we identified one candidate (NY1-B04), which was >2 log_2_FC enriched when comparing sorted cells to cells prior to sorting (Figure 3b-c). NY1-B04 pMHC binding characteristics aligned with the MD predictions and demonstrated the utility of this approach to deselect false positive binders. NY1-B04 was individually cloned, and binding of SLLMWITQC/HLA-A*02:01 tetramers were confirmed in CD3 KO Jurkat cells (Figure 3d). To investigate the cross-reactivity profile of NY1-B04, we performed a cross-reactivity experiment by staining the NY1-B04^+^ CD3 KO Jurkat cells with selected pMHC tetramers. We tested the ability of NY1-B04 to bind 9 of the 10 variant peptides tested in the *in silico* cross-panning in complex with HLA-A*02:01 (Figure 3e-g, Supplementary Figure S2). In line with the *in silico* cross-panning predictions, NY1-B04 was cross-binding to the single mutants SLL**A**WITQC and SLLMWITQ**V**, which were also predicted to bind in the *in silico* cross-panning. NY1-B04 also bound SLL**EH**ITQC, however not to the same extent as the original SLLMWITQC peptide and SLLAWITQC and SLLMWITQV variants (Figure 3e-g). This suggests the identified binder possesses limited cross-binding to similar peptide sequences and that the *in silico* cross-panning is useful (accuracy_N=9_ = 8/9 peptide variants, 1 false negative in silico prediction) for identifying cross-reactive peptides for true binders (Supplementary Figure S2).

**Figure 3.**
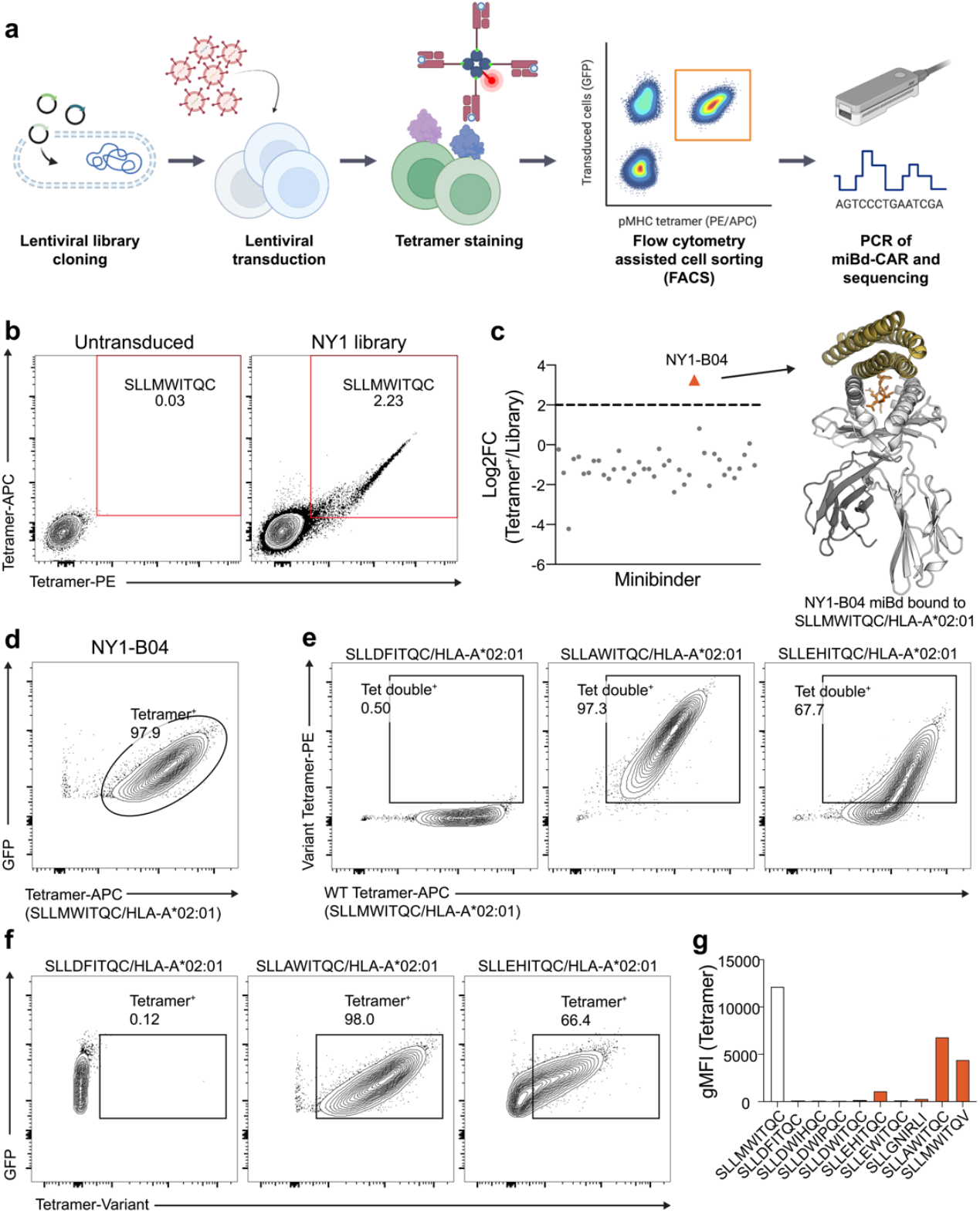
Experimental selection and characterisation of binders. **(a)** Mammalian cell display workflow. **(b-d)** A library of 44 *de-novo* generated miBds was lentivirally transduced into CD3 KO Jurkat cells. Cells were stained with SLLMWITQC/HLA-A*02:01 tetramers (PE and APC), double-positive cells were sorted and the integrated miBds were sequenced. **(b)** Flow plots showing the sorted cells and **(c)** log2FC of miBds in the sorted tetramer^+^ cells compared to the cells prior to sorting for the identification of NY1-B04 miBd targeting SLLMWITQC/HLA-A*02:01. **(d)** NY1-B04 binding to SLLMWITQC/HLA-A*02:01 was confirmed by a single tetramer-APC stain. **(e-f)** Cross-reactivity screening of NY1-B04 against variant peptides showing 3 examples of non-binding and binding peptides following double staining with SLLMWITQC/HLA-A*02:01 and variant tetramers (e) or single tetramer staining with the variant tetramers (f). **(g)** Bar plot showing geometric mean fluorescence intensity (gMFI) of the single-stained tetramer stains. Gated on live CD3^-^ GFP^+^ cells in (b), (d-f).

### Killing cancer cells with miBd-CAR T cells

To evaluate the functional potential of NY1-B04 as a cancer-targeting molecule, and not just target binding, we tested if the ‘NY1-B04’-miBd could induce killing in the context of a BIKE. We generated primary human T cells expressing an ‘NY1-B04’-CD8α hinge-CD28-CD3ζ CAR (BIKE) by lentiviral transduction (Figure 4a,b). Subsequently, we evaluated their capacity to kill NY-ESO-1^+^ A375 melanoma cells (expressing mCherry as a reporter). We demonstrated that the NY1-B04 BIKE-T cells induced rapid cell death of the A375 cell line within 24h (Figure 4c). Apoptotic bodies of A375 cells were clearly visible after 24h, and the A375 confluence rapidly declined. A375 cell death was significantly higher in BIKE conditions compared to the non-transduced control, providing a first proof of principle that these *de novo-*designed miBds have the potential to play a significant role in cell-based therapies.

**Figure 4.**
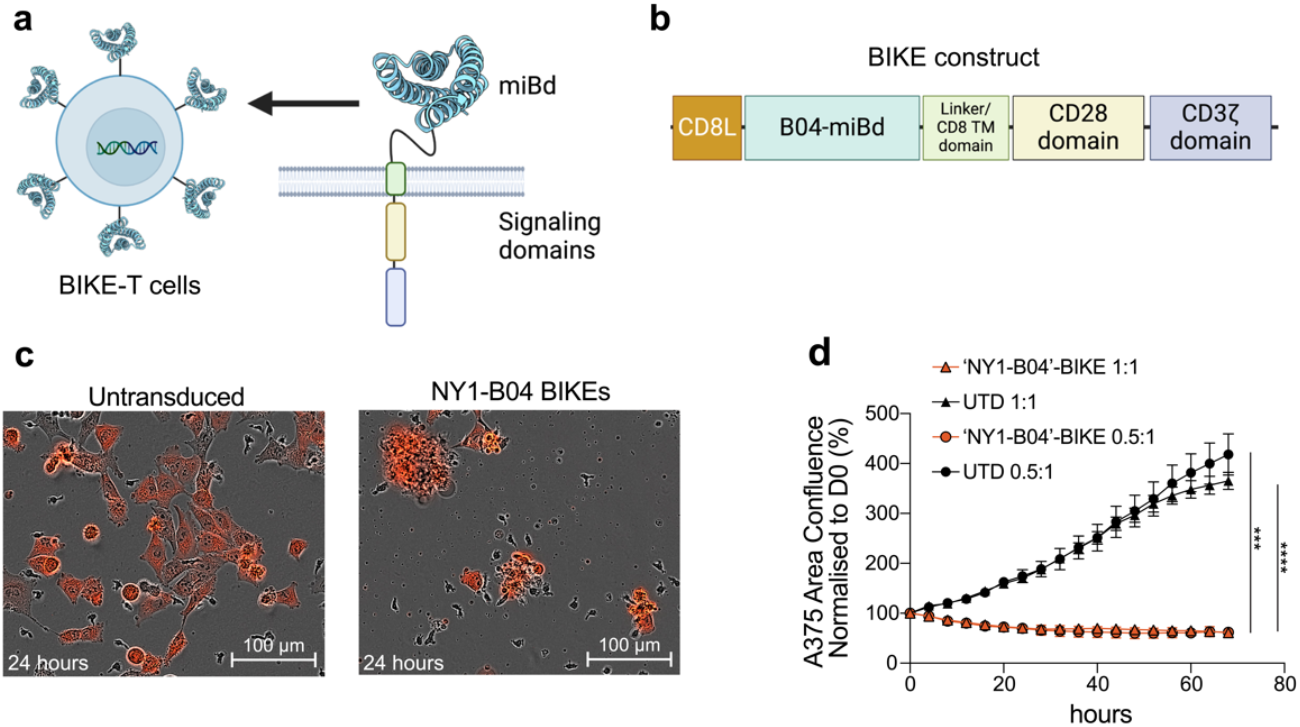
BIKE T cell mediated cytotoxicity against NY-ESO-1^+^ A375 cancer cells. **(a)** Overview of the *de novo* pMHC-Binder for Immune-mediated Killing Engager (BIKE) T cells concept with miBds fused extracellularly to intracellular signalling domains. **(b)** Construct design of the ‘NY1-B04’-BIKE with a CD8 leader, miBd, CD8a hinge region, and CD28/CD3ζ intracellular signalling domains. **(c)** Phase contrast images overlaid with mCherry fluorescence showing the mCherry expressing A375 cancer cells after 24 hours of incubation of non-transduced (UTD) primary T cells (left) or ‘NY1-B04’-BIKE-transduced primary T cells (right). **(d)** Cell death of mCherry-expressing A375 cancer cells was monitored and quantified over a period of ∼2 days using Incucyte® live-cell imaging, with red fluorescence area normalised to D0 area as a measure of confluency. The assay was set up with 0.5:1 and 1:1 effector:target ratio. P≤0.001:***, P≤0.0001:**** as evaluated by unpaired t-test comparing the area confluence of the last timepoint between UTD and ‘NY1-B04’-BIKE samples.

### Extending the miBd-discovery pipeline to neoantigens

A key hurdle limiting the utility of this technology would be the reliance on experimentally solved and extensively characterised pMHC targets^2^, such as SLLMWITQC/HLA-A*02:01. We, therefore, sought to further test the limits of our approach by targeting the neoantigen RVTDESILSY/HLA-A*01:01. The metastatic melanoma neoantigen, RVTDESILSY/HLA-A*01:01 derived from the mutation AKAP9^P1796L^, was previously identified based on T-cell recognition in both blood and tumour infiltrating lymphocyte (TIL) infusion product^22^. This neoepitope is considered private, not being broadly reported in melanoma patients, and is derived from a passenger mutation that does not drive tumour growth. Based on structure predictions, the neoantigen results in a targetable protruding leucine in the MHC-I-groove^22^. Since no experimental structure existed we proceeded by modelling the pMHC complex using AF2. Upon selecting the most confident structure based on AF2 (pLDDT of 95.7), we designed miBds as described above (Figure 5a) and observed similar structural characteristics as for the SLLMWITQC/HLA-A*02:01 miBds (Extended Data Table 2). We selected 95 miBds for *in vitro* testing, and we found that the fluorochrome intensity of the tetramer^+^ miBd library was comparable to that of primary T cells with CRISPR/Cas9 knockin of an RVTDESILSY/HLA-A*01:01-specific TCR originally identified in the patient, as evidenced by a gMFI of 14103 (TCR) and 18546 (miBd) (Figure 5b). Sequencing of the sorted population identified two unique binders (i.e., SILSY1-B11 and SILSY1-G05), constituting a success rate of ∼2% (Figure 5c). Importantly, this demonstrated the possibility of applying this design pipeline to neoantigens on other HLAs than HLA-A*02:01 and targets with no experimental structure available, for example, in the context of personalised cancer therapy.

**Figure 5.**
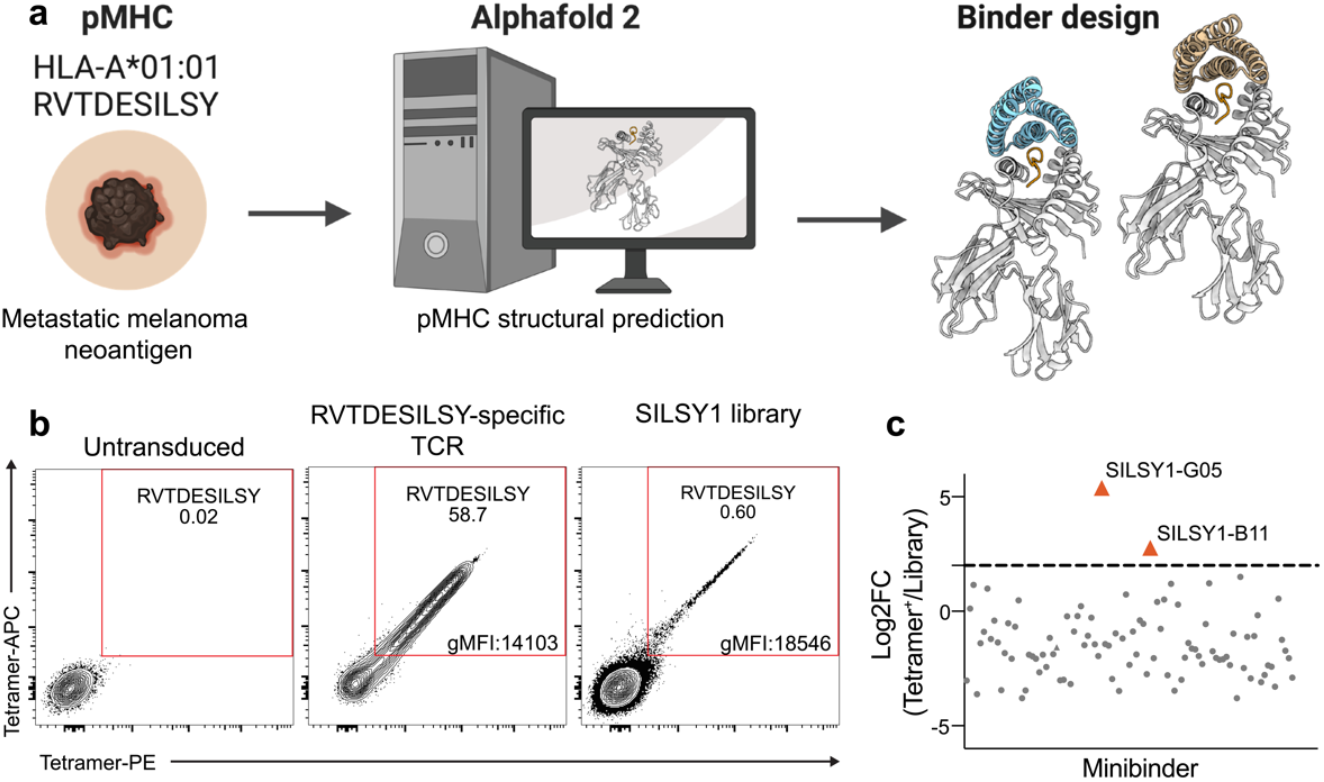
Proof of concept of applying the design pipeline to neoantigens. **(a)** The structure of the neoantigen RVTDESILSY in complex with HLA-A*01:01 was predicted using AF2 and miBds designed following the previously described pipeline. (b) A library of 95 predicted miBds was lentivirally transduced in CD3 KO Jurkat cells and cells were stained with RVTDESILSY/HLA-A*01:01 tetramers (PE and APC) and double positive populations were sorted and the integrated miBds were sequenced. **(c)** Analysing the log2FC of miBds represented in the sorted population (Tetramer^+^) versus the cells prior to sorting (Library) identified two miBds (SILSY1-B11 and SILSY1-G05 in red) targeting RVTDESILSY/HLA-A*01:01. Log_2_FC of 2 is marked with a dotted line.

## DISCUSSION

In cancer immunotherapy a significant challenge has been pMHC-targeting with high specificity and efficacy^23–25^. Traditionally this has been achieved using natural TCRs identified by screening T-cell responsiveness and specificity from cancer patients^26–30^. This, however, is extremely time and resource-intensive while also often lacking the required specificity, potentially not just preventing patient treatment but also with the risk of fatal side effects^31^. Thus, there has been a major push toward novel approaches that can help overcome these challenges and help unlock the huge potential that targeted T-cell therapy holds. Here, we approached pMHC-targeting by leveraging leading-edge protein design approaches to rapidly generatively design anti-SLLMWITQC/HLA-A*02:01 pMHC binders *de novo*^16^. We further enhanced current design pipelines employed by us and others^16,32,33^ to screen for specificity *in silico* and reduce the risk of off-target pMHC. We demonstrated that these rapid (<10 sec per miBd-target complex) computational predictions correlated substantially (89%) with *in vitro* specificity screening results, substantiating the potential this approach has in de-risking T-cell therapy. This specificity then also resulted in highly effective and specific target cell killing when using the miBd as the recognition element in a CAR; this novel type of pMHC recognition receptor (BIKEs) can direct T cells towards the pMHC antigen of interest, resulting in receptor engagement, signalling and consequently cytotoxicity towards the pMHC-expressing target cell. Peptide-centric CARs that target pMHCs have been tested and shown success *in vitro* and *in vivo*^7^; however, our BIKE approach allows for more rapid discovery with the added benefit of the implemented *in silico* cross-reactivity screening. This holds great promise for a rapid therapeutic discovery pipeline to target both shared cancer antigens such as NY-ESO-1, but also patient-specific pMHC targets such as neoantigens^34^, since one could go from design to therapeutic cell product based on patient T cells transduced with BIKEs in under 8 weeks. Another major strength of this approach is the relatively high success rate (1:50) of identification of novel binders, which we demonstrated to be able to further increase using molecular dynamics approaches. We found that higher variability in the protein interface and higher fluctuations in interaction energies were important features to deselect potential pMHC-miBds - a comparison facilitated by the high similarity of the binding interfaces among these potential binders. Thus, we propose that considering interface flexibility and dynamics can substantially reduce the false positive rate and further expedite the identification of the most promising miBd-pMHC complex.

The presented work and workflow constitute a major leap forward in T-cell therapy that could guide the way for a plethora of cancer therapies. Beyond cancer applications, the versatility of miBds makes them promising for targeting viral antigens presented on infected cells. This could be transformative for diseases such as HIV or hepatitis C, where broad-spectrum targeting remains challenging due to high mutation rates. Additionally, miBds could be used to block autoimmune TCR-pMHC interactions, as proposed for TCR-like antibodies^35^, further expanding their therapeutic scope. Further, the findings shown here unlock the use of pMHC-binding miBds as carriers of payloads or pMHC-targeting bispecific T cell engagers (biTes). Given the previous success of generatively designed binders *in vivo*^33^ these findings are likely to not only advance the field of cancer immunotherapy by introducing a rapid and precise method for *de novo* pMHC targeting, but also pave the way for targeted patient-centric and, if required, personalised therapies.

## METHODS

### *De novo* pMHC binder design using RFdiffusion

The crystal structure of NY-ESO-1, SLLMWITQC/HLA-A*02:01 (PDB ID: 2BNR) was used as input to RFdiffusion. 5,500 diffused designs were generated using RFdiffusion^16^, with all amino acids from the peptide serving as hotspots to increase the chance of binding being primarily driven by the peptide rather than the MHC. For each backbone structure, four amino acid sequences of putative pMHC binder were designed using ProteinMPNN^17^ and scored via AF2 with *initial guess*^18,19^. The binders were filtered based on interaction predicted alignment error (ipAE) of the pMHC:binder complex and the predicted local distance difference test (pLDDT). All binders with an ipAE score below 12 Å and a pLDDT score above 88 were selected for further optimisation. The subpopulation of selected designs surpassing the ipAE and binder pLDDT were further filtered for unique initial RFdiffusion backbones to maximise the structural diversity; this resulted in a set of 109 binder designs (Table 1). Following the initial selection, the designed binders were subjected to sequence diversification. Since the initial design and selection process resulted in a high structural variety of >100 binder designs for the pMHC SLLMWITQC/HLA-A*02:01, only sequence and no structural diversification were performed. At the default sampling temperature t = 0.1, 100 sequences were generated per initial binder design and reinterpreted into 10,000 structures by AF2, whereafter, the ipAE and pLDDT were extracted as described above. The best scoring variant for each initial binder design was identified, and the subpopulation was filtered based on an ipAE cut-off of 7 Å and a pLDDT cut-off of 92, which resulted in a subset of 44 pMHC binders.

**Table 1.** Overview of binder design strategy against NY-ESO-1.

| Target | Best binder (ipAE) [Å] | Number of initial backbone designs | Hot spot selection | ProteinMPNN sequence designs & AF2 | Selected designs after initial round | Refinement: ProteinMPNN & AF2 re-interpretation | Final set of selected designs for in vitro testing |
| --- | --- | --- | --- | --- | --- | --- | --- |
| NY-ESO-1: SLLMWITQC/HLA-A*02:01 | 4.5 | 5500 | All peptide positions: C1-C9 | 4 sequences per 5500 structural designs | 109 | ProteinMPNN & AF2 re-interpretation (100 sequences per design) | 44 |

### *In silico* cross-panning in AF2

The selected 44 pMHC binders for the SLLMWITQC/HLA-A*02:01 complexes were evaluated for their predicted binding capabilities against the MHC containing no or mutated variants of the target peptide to attempt decreasing the chance of off-target binding. First, all 3,610 conceivable single and double-point mutated peptides were generated (while keeping peptide positions 1-3 and 9 unchanged, as described above). These 3,610 peptides were blasted against the human proteome using NCBI pBlast^36^, which led to the identification of 16 hits with a length between 5 and 10 amino acids (at a 85% identity) - the predominant match was the target peptide itself (Supplementary Figure S1). The MHC affinity of the 16 peptides was predicted using NetMHCpan4.1^37^, and only two peptides (SLLGNILRI and SLLGNILRII) scored equally well or better than the original peptide (SLLMWITQC). Due to the significant similarity of these two peptides only SLLGNILRI was included in the following cross-reactivity screening. As a second strategy of cross-reactivity screening, all 3,610 peptides were also subjected to MHC affinity prediction using NetMHCpan4.1. Of the 3,610 peptides, 1,128 scored higher than the original peptide, indicating these were more likely to be presented on the HLA-A*02:01 complex. Overlaying the sequences in a logo plot it became apparent the residue in position 4, and to a lesser extent also the residues in positions 5, 7, and 8 were the least conserved residues with no or only a marginally dominating amino acid (Supplementary Figure S2). To reduce the requirement for computational resources, only 10 single- or double-point mutated peptides were chosen for structural modelling in AF3 and subsequent cross-reactivity screening with AF2, resulting in a total combination of 484 structures: (1 peptide from strategy A, 9 mutated peptides from strategy B, 1 remodelled original peptide) x 1 HLA-A*02:01 complex x 44 selected pMHC binders. Upon remodelling and scoring the pMHC binders using AF2 with *initial guess*, none of the miBds were predicted to be only binding to the original SLLMWITQC peptide, but instead likewise obtained obtained low ipAE scores (<10) towards the peptide variants SLLAWITQC/HLA-A*02:01 and SLLMWITQV/HLA-A*02:01 (Figure 1c). Lastly, not all miBd designs scored below ipAE < 10 against the original, but remodelled version (by AF3) of SLLMWITQC/HLA-A*02:01. All 44 designs were, nevertheless, selected for further *in vitro* validation.

### *In silico* structural characterisation of the binders

To quantitatively assess the secondary structure configuration, we assessed the ψ and ϕ angle between the protein backbone nitrogen atom and the carbon α atom for the first and between the carbon α atom and the carbon atom involved in the carboxyl group for each amino acid from all miBd-pMHC complexes, for the latter angle (Extended Data Table 1). Utilising the characteristic angle ranges for ψ and ϕ residues were classified into zones defining ‘β sheets’, ‘right-handed α helices’ and ‘left-handed α helices’ (Extended Data Table 1). For all pMHC binders, independent of the design campaign and the peptide they have been designed against right-handed α helices were dominant in the secondary structure. Generally, about 91-97% of the amino acids in the pMHC binder were located in right-handed α helices (Table 2). The principal axes and geometric centers were calculated using singular value decomposition on the α carbon coordinates of each helix and the peptide. An overall orientation label of the miBds was determined by averaging the angles of all helices relative to the peptide’s principal axis. Angles greater than 90° were adjusted in order to treat parallel and anti-parallel alignments as equivalent. A logistic regression model was implemented to classify miBd orientations. The model incorporated features such as helix-peptide angles and distances. L2 regularisation was applied and regularisation strength (λ) was optimised using 10-fold cross-validation, where the optimal λ value was selected based on the lowest cross-validation error. Next, the orientation of the C-and N-termini of the pMHC binders was investigated by measuring the distance between the carbon α atoms of each terminus as well as the distance to the individual positions of the peptides. If the distance between the C- or N-terminus towards the peptide positions increased from the peptide’s C-to N-terminus, the orientation was classified as ‘dorsal’. If the distances decreased, the orientation would be classified as ‘frontal’. Lastly, if the distances led to indifferent conclusions, the orientation was classified as ‘lateral’, which entails every configuration of the C- and N-termini that is not strictly perpendicular to the peptide.

**Table 2.** Overview of binder design strategy against RVTDESILSY.

| Target | Best binder (ipAE) [Å] | Number of initial backbone designs | Hot spot selection | ProteinMPNN sequence designs & AF2 in initial round | Selected designs after initial round | Refinement: ProteinMPNN & AF2 re-interpretation | Final set of selected designs for in vitro testing |
| --- | --- | --- | --- | --- | --- | --- | --- |
| RVTDESILSY/HLA-A*01:01 | 5.2 (initial selected designs)<br>3.4 (Strategy 2: partial diffusion)<br>3.2 (Strategy 1) | 2100 (initial campaign) | All peptide positions: C1-C10<br>Non-Anchor peptide residue positions: C3-C7<br>Mutated residue position: C7 | 4 sequences per 2100 structural designs | 30 for ProteinMPNN | Strategy 1: ProteinMPNN & AF2 reinterpretation (sampling temperature $t=0.1$ and $N=30$ sequences per initial design)<br>Strategy 2: Partial diffusion (with 20 timesteps and $n=4$ sequences per partially diffused structure) | 96 |

### Molecular dynamics simulations and analysis

Using the structure models from AF2 as starting structures, we performed molecular dynamics simulations of the de-novo-designed miBds to identify descriptors that can facilitate the downselection of miBds to reduce the rate of false positives. The starting structures for our simulations were prepared in MOE^38^ using the Protonate3D tool^39^. To neutralise the charges, the uniform background charge was applied, which is required to compute long-range electrostatic interactions^40^. Using the tleap tool of the AmberTools^41^ package, the structures were soaked in cubic water boxes of TIP3P water molecules with a minimum wall distance of 12 Å to the protein^42–44^. For all simulations, parameters of the AMBER force field 14SB were used^45^.

We performed each 4 × 500 ns of classical molecular dynamics simulations using the AMBER 21 simulation software package, which contains the GPU implementation of the Particle Mesh Ewald MD method (pmemd.cuda). Bonds involving hydrogen atoms were restrained using the SHAKE algorithm, allowing a timestep of 2.0 femtoseconds. The Langevin thermostat was used to maintain the temperature during simulations at 300 K^46,47^ with a collision frequency of 2 ps−1 and a Monte Carlo barostat_48_ with one volume change attempt per 100 steps.

The interaction energies and the respective fluctuations were calculated with cpptraj using the interaction energy (LIE) tool^41,49^. The electrostatic and Van der Waals interaction energies were calculated for all frames of each simulation, and the fluctuations of the energies were used as parameters to facilitate selection of true positives (Figure 2). To calculate the interactions and interaction frequencies of the miBd-pMHC binding interfaces over the simulations, we used the GetContacts tool (Stanford University, adate; https://getcontacts.github.io/). The interface residues forming salt-bridge and hydrogen bond interactions for more than 10% of the simulation are considered for calculating an interface-residue B-factor, as an estimator for interface flexibility and stability. The B-factor was calculated with cpptraj^49^. The radar plot combining all three descriptors (individually normalised) to characterise protein interface flexibility and variability was generated with NumPy^50^ and Matplotlib^51^. PyMOL was used for visualising protein structures (The PyMOL Molecular Graphics System, Version 2.5.2 Schrödinger, LLC).

### Recombinant expression of MHC molecules and generation of pMHC complexes

MHC molecules were expressed as previously described^52^. In brief, recombinant HLA-A*02:01 heavy chain carrying a Y84C mutation, HLA-A*01:01 heavy chain and human β_2_ microglobulin light chain were produced in *Escherichia coli*. HLA-A*02:01 heavy and light chains were refolded with 10 mM dipeptide GM, while HLA-A*01:01 heavy and light chains were refolded with 60 uM of photocleavable peptide KILGFVF-J-V. All proteins were then biotinylated and purified by size exclusion chromatography using high-performance liquid chromatography (Waters Corporation, USA). For generating specific peptide-HLA-A*02:01-Y84C, purified HLA-A*02:01-Y84C monomers (200 μg/mL) were incubated with 400 uM peptide in phosphate-buffered saline (PBS) for 30 min at RT. RVTDESILSY/HLA-A*01:01 were generated by UV-mediated peptide exchange by incubating purified HLA-A*01:01 monomers (200 μg/mL) with 400 uM peptide in phosphate-buffered saline (PBS) for 1 hr at RT under UV light (366 nm). Specific pMHC complexes were centrifuged for 5 min at 10’000 *g*, and supernatants were used to generate pMHC tetramers.

### Preparation of pMHC tetramers

For each 100 ul of pMHC monomers, 9 ul of [0.2 mg/mL stock, SA-PE (Biolegend, Cat.#405204), and SA-APC (Biolegend, Cat.#405243)] of streptavidin conjugates was added and incubated for 30 min at 4°C, followed by the addition of D-biotin (Sigma-Aldrich) at a final concentration of 25 uM to block any free binding sites. pMHC tetramers were stored at -20°C with 5% glycerol and 0.5% BSA.

### Maintenance and modification of cell lines

The A375 melanoma cell line (kindly gifted by Dr. Marco Donia, CCIT), HEK293T cells (ATCC, 293T Cat.#CRL-3216), and the Jurkat cell lines (Signosis, Cat.#SL-0032) were cultured in complete R10 (ATCC modification, Thermo Fisher, Cat.#A1049101) [RPMI 1640 + 10 % FBS + 1% Pen-Strep] at 37°C, 5 % CO2, and split every 2-4 days before confluence.

A375 cells were modified to express mCherry and Luciferase by lentiviral transduction of pDTU_mCherry_Luciferase (Supplementary Table S3) and sorted based on bright mCherry expression on an Aria Fusion (BD Bioscience) to generate mCherry^+^ A375 cells. To produce Jurkat CD3 KO cells, Jurkat cells were electroporated with Ribonucleoproteins (RNPs) targeting CD3ε (CD3ε crRNA in Supplementary Table S3). The resulting CD3-negative population was sorted in bulk after 5 days. All cell lines were regularly tested for Mycoplasma by qPCR.

### Design and cloning of genes

MiBd libraries of 44 designs against SLLMWITQC/HLA-A*02:01 and a 95 design library against RVTDESILSY/HLA-A*01:01 were synthesised as gBlocks™ Gene Fragment (Integrated DNA Technologies) or as Synthetic Gene Fragments (Twist Bioscience). The two libraries were cloned as separate pools into the CAR transfer vector (pMB-LINK), a modified version of pLenti-puro (Addgene, #39481), optimised to include a cPPT-CTS sequence, the EF-1α promoter, a Woodchuck Hepatitis Virus (WHV) Post-transcriptional Regulatory Element (WPRE), a CD8 leader sequence, BsmBI cloning sites, the CAR domains CD8α hinge-CD28-CD3ζ, and GFP^53^ (Supplementary Table S3).

Cloning of miBds into pMB-LINK was performed using NEBridge® Golden Gate Assembly Kit (BsmBI-v2) (NEB) according to manufacturer’s instructions. Assemblies were electroporated into ElectroMAX STBL4 competent bacteria with the program ‘*E. coli* – 1mm,1.8 kV’ on Gene Pulser Xcell Electroporation System (BioRad). Library coverage was assessed by PCR using NEBNext® Ultra™ II Q5® Master Mix (NEB) and primers MB_GGinterm_fwd_2 and MB_GGinterm_rev_2 (Supplementary Table S3). Amplified miBd sequences were labelled with Native Barcoding Kit 96 V14 (SQK-NBD114.96, Nanopore) and sequenced by Nanopore MinIon. For data analysis, FASTQ read files obtained after sequencing were trimmed using a custom Python script and matched against the library of miBd sequences.

### Mammalian miBd display screening

Lentiviral particles containing the miBds libraries were produced by lipofectamine-based co-transfection of HEK293 cells with 3rd generation packaging plasmids pMD2.G (Addgene, Cat.#12259), pMDLg/pRRE (Addgene, Cat.#12251), pRSV-Rev (Addgene, Cat.#12253), and the pMB-LINK. 24 and 48 hrs hrs after transfection, lentiviral particles were harvested and concentrated using Lenti-X concentrator (Takara Bio) and stored at -80°C. CD3 KO Jurkat cell lines were transduced with lentivirus at an MOI of 1 to ensure one integration per cell.

Tetramers were separately labelled with PE and APC fluorochromes and pooled before cell staining. For NY-ESO-1-binders, SLLMWITQC/HLA-A*02:01 were used, and for SILSY-binders, RVTDESILSY/HLA-A*01:01 tetramers were used. CD3 KO Jurkat cell lines were stained with mouse anti-Human CD3 (BD, #562877) and LIVE/DEAD™ Fixable Near-IR Dead Cell Stain Kit (Thermofisher scientific). Binder expression was assessed by evaluating the GFP signal, while pMHC binding to the surface-expressed pMHC binder was confirmed by double PE- and APC-tetramer staining. Cells were sorted for CD3^-^, GFP^+^ and double positive for PE and APC on a BD FACSDiscover™ S8 Cell Sorter (BD, USA). Genomic DNA was extracted using QIAamp DNA Micro Kit (Qiagen) following manufacturer’s instructions. To amplify the integrated miBd DNA from the integrated CAR construct, PCR and sequencing were performed as described before. FASTQ read files were analysed as outlined above. Read counts for each miBd were normalised against total read counts in cells before sorting. For each miBd, the log2-transformed fold change (log_2_FC) for enrichment was calculated between the sorted samples and the representation of miBds before sorting. The identified binder sequences for SLLMWITQC/HLA-A*02:01 were cloned as single constructs and transduced by lentiviral transduction as described in ‘Design and cloning of genes’ to confirm binding. To evaluate specificity, cross-reactivity screens were performed by separately staining the transduced CD3 KO Jurkat cell lines with the target SLLMWITQC/HLA-A*02:01 tetramers in one fluorophore (PE) and a range of undesired pMHC targets tetramers in another fluorophore (APC). In particular, the undesired pMHC tetramers encompassed single-, double-point or quadruple-point mutated peptides bound to HLA-A*02:01 (Supplementary Table S3).

### Incucyte killing assays

The identified NY1-B04 binder was cloned into the pMB-LINK vector used for mammalian display. Primary T cells were isolated from peripheral blood mononuclear cells (PBMC) from healthy donors and activated overnight with 5 μg/mL plate-bound anti-CD3, 2 μg/mL anti-CD28 and 20 IU/mL of Interleukin-2 (IL-2). The following day, activated T cells were transduced with the ‘NY1-B04’-CD8α-CD28-CD3ζ BIKE construct. Primary human T cells were cultured in X-vivo-15 medium (Lonza) with 5% human serum and 1% pen-strep (X-vivo-15+5%HS+1%PS) at 37°C with 5% CO_2_. A375 cells were seeded in flat-bottom 96-wells the day before the addition of T cells. At day 10, transduced primary T cells were mixed 0.5:1 and 1:1 with mCherry^+^ labelled A375 cancer cells. The cytotoxicity of the ‘NY1-B04’-BIKE T cells was evaluated using an Incucyte Live Cell Analyzer (Sartorius), which took images every 4 hours over 72 hours, quantifying the number of live target cells over time. Data was analysed in Prism 10.4.0 (Graphpad), and the last time point was compared between UTD and NY1-B04 transduced cells by unpaired students *t*-test.

### Discovery of neoantigen binders

The structure of the neoantigen pMHC complex, RVTDESILSY/HLA-A*01:01, was predicted using the ColabFold implementation of AF2^18,54^ with standard settings. The input included the MHC allele sequence and the neoantigen peptide, ensuring accurate modelling of the peptide binding groove and structural conformation. We selected the most confident model and proceeded designing miBds as described above (Table 2).

We then selected the 30 best scoring (ipAE and pLDDT) RVTDESILSY/HLA-A*01:01 miBds for further *in silico* diversification. We employed two strategies: sequence only, as well as sequence and backbone diversification. In the former, the backbones of the 30 selected miBds were re-interpreted using ProteinMPNN at default sampling temperature t = 0.1 and N = 30 sequences per initial binder design.

In the second strategy, all 30 selected pMHC binder designs were re-interpreted structurally via partial diffusion with denoising steps T = 20 and, subsequently, in sequence with N = 4 sequences per partially diffused structure binder design. Next, the newly generated variants of the 30 selected pMHC binder designs from both strategies were filtered based on an ipAE < 7 Å and pLDDT > 80 %. 1117 pMHC binder designs passed the threshold, of which 413 originated from the sequence only, while 704 came from the backbone and sequence strategy. A further 127 of the 1117 pMHC binder designs revealed clash reports when analysed by TopModel^55^ as described above. Notably, two of the designs initial 30 best-scoring designs were also reported to have several clashes, which resulted in their deselection. In total, 68 pMHC binder designs were selected for *in vitro* screening alongside 28 of the original 30 pMHC miBds (n=96). Similarly, as above, we quantitatively assessed the secondary structure configuration of all the selected miBd-pMHC complexes, respectively (Extended Data Table 2).

## Code availability

Code explanation and examples of binder design using RFdiffusion can be found at https://github.com/RosettaCommons/RFdiffusion#binder-design

The Pymol script for assessment and depiction of potential clashes of co-complexes is available on GitHub by accessing GitHub - liedllab/TopModel: A structure checker in python

## Acknowledgements

T.P.J. acknowledges support from the Alliance programme under the EuroTech Universities agreement. D.S.W. is very thankful for his PhD fellowship stipend granted by Innovation Fund Denmark [2052– 00010B (2022)]. KHJ was kindly funded by a Lundbeckfonden grant [R347-2020-2174] SRH is a grateful recipient of a NNF project grant [0087077] and ERC consolidator grant MIMIC, both supporting the project. MLFQ acknowledges EuroHPC Joint Undertaking for awarding us access to MeluXina, Luxemburg. We thank Carlos Rodriquez Pardo for CD3 KO Jurkats cells and Sebastian Sebbaha from the FLIC core facility for supporting flow cytometry experimental design and data analysis. We also thank Bingxu Liu, Nathan F. Greenwood, and Julia Bonzanini from the Baker lab for discussions and knowledge exchange, as well as for timing submissions to BioRxiv.

## Author Contributions

T.P.J., K.H.J, and S.R.H conceptualised the project. T.P.J., K.H.J, and S.R.H provided direction for the work. D.S.W. designed and *in silico* screened the pMHC binders with the help of C.R.C. and T.P.J.. M.D. and A.T.D.E. initiated the design of neoantigen pMHC binder together with D.S.W. and C.R.C.. E.Q.A. and D.S.W. analysed the secondary structure composition and orientation of the miBds. K.H.J, B.S., and C.R.C. prepared the binder libraries, screened via mammalian display and sequenced the hits. M.F.Q, J.R.L, S.F. and A.B.W. conducted all MD experiments. K.H.J., B.S, and C.R.C. conducted all cell assays. K.K.M. analysed the predicted miBd structures and aided with sequencing and data analysis. M.D.O., O. M., and D.S.W. initiated and improved the protocols for *in silico* specificity screening. M.O. assisted with CAR vector design. T.P.J., K.H.J., B.S., D.S.W., and S.R.H. wrote the original draft of the manuscript. All authors reviewed and accepted the manuscript.

## Requests for materials

Correspondence and requests for materials should be addressed to T.P.J., K.H.J., and S.R.H.

## References

1. Pétremand, R. et al. Identification of clinically relevant T cell receptors for personalized T cell therapy using combinatorial algorithms. Nat. Biotechnol. 1–6 (2024) doi:10.1038/s41587-024-02232-0.

2. Arnaud, M. et al. Sensitive identification of neoantigens and cognate TCRs in human solid tumors. Nat. Biotechnol. 40, 656 (2021).

3. Strønen, E. et al. Targeting of cancer neoantigens with donor-derived T cell receptor repertoires. Science (2016) doi:10.1126/science.aaf2288.

4. Drost, F. et al. Predicting T cell receptor functionality against mutant epitopes. Cell Genomics 4, 100634 (2024).

5. Nielsen, M. et al. Lessons learned from the IMMREP23 TCR-epitope prediction challenge. ImmunoInformatics 16, 100045 (2024).

6. Klebanoff, C. A., Chandran, S. S., Baker, B. M., Quezada, S. A. & Ribas, A. T cell receptor therapeutics: immunological targeting of the intracellular cancer proteome. Nat. Rev. Drug Discov. 22, 996–1017 (2023).

7. Yarmarkovich, M. et al. Targeting of intracellular oncoproteins with peptide-centric CARs. Nature 623, 820–827 (2023).

8. Dao, T., Liu, C. & Scheinberg, D. A. Approaching untargetable tumor-associated antigens with antibodies. OncoImmunology 2, e24678 (2013).

9. Dao, T. et al. Targeting the Intracellular WT1 Oncogene Product with a Therapeutic Human Antibody. Sci. Transl. Med. 5, (2013).

10. Li, Y., Jiang, W. & Mellins, E. D. TCR-like antibodies targeting autoantigen-mhc complexes: a mini-review. Front. Immunol. 13, 968432 (2022).

11. Du, H. et al. A general platform for targeting MHC-II antigens via a single loop. bioRxiv (2024).

12. Stewart-Jones, G. et al. Rational development of high-affinity T-cell receptor-like antibodies. Proc. Natl. Acad. Sci. 106, 5784–5788 (2009).

13. Lee, C. H. et al. Predicting cross-reactivity and antigen specificity of T cell receptors. Front. Immunol. 11, 565096 (2020).

14. Karnaukhov, V. K. et al. Structure-based prediction of T cell receptor recognition of unseen epitopes using TCRen. Nat. Comput. Sci. 4, 510–521 (2024).

15. Zhao, Y. et al. DeepAIR: A deep learning framework for effective integration of sequence and 3D structure to enable adaptive immune receptor analysis. Sci. Adv. 9, eabo5128 (2023).

16. Watson, J. L. et al. De novo design of protein structure and function with RFdiffusion. Nature 620, 1089–1100 (2023).

17. Dauparas, J. et al. Robust deep learning–based protein sequence design using ProteinMPNN. Science 378, 49–56 (2022).

18. Jumper, J. et al. Highly accurate protein structure prediction with AlphaFold. Nature 596, 583–589 (2021).

19. Jumper, J. & Hassabis, D. Protein structure predictions to atomic accuracy with AlphaFold. Nat. Methods 19, 11–12 (2022).

20. Chen, Y.-T. et al. A testicular antigen aberrantly expressed in human cancers detected by autologous antibody screening. Proc. Natl. Acad. Sci. 94, 1914–1918 (1997).

21. Abramson, J. et al. Accurate structure prediction of biomolecular interactions with AlphaFold 3. Nature 630, 493–500 (2024).

22. Kristensen, N. P. et al. Neoantigen-reactive CD8^+^ T cells affect clinical outcome of adoptive cell therapy with tumor-infiltrating lymphocytes in melanoma. J. Clin. Invest. 132, (2022).

23. Johnson, L. A. et al. Gene therapy with human and mouse T-cell receptors mediates cancer regression and targets normal tissues expressing cognate antigen. Blood 114, 535–546 (2009).

24. Linette, G. P. et al. Cardiovascular toxicity and titin cross-reactivity of affinity-enhanced T cells in myeloma and melanoma. Blood 122, 863–871 (2013).

25. Parkhurst, M. R. et al. T Cells Targeting Carcinoembryonic Antigen Can Mediate Regression of Metastatic Colorectal Cancer but Induce Severe Transient Colitis. Mol. Ther. 19, 620–626 (2011).

26. Morgan, R. A. et al. Cancer Regression in Patients After Transfer of Genetically Engineered Lymphocytes. Science 314, 126–129 (2006).

27. Klebanoff, C. A., Chandran, S. S., Baker, B. M., Quezada, S. A. & Ribas, A. T cell receptor therapeutics: immunologic targeting of the intracellular cancer proteome. Nat. Rev. Drug Discov. 22, 996–1017 (2023).

28. Chour, W. et al. Large libraries of single-chain trimer peptide-MHCs enable antigen-specific CD8+ T cell discovery and analysis. Commun. Biol. 6, 1–13 (2023).

29. Moravec, Z. et al. Discovery of tumor-reactive T cell receptors by massively parallel library synthesis and screening. Nat. Biotechnol. (2024) doi:10.1038/s41587-024-02210-6.

30. Giannakopoulou, E. et al. A T cell receptor targeting a recurrent driver mutation in FLT3 mediates elimination of primary human acute myeloid leukemia in vivo. Nat. Cancer 4, 1474–1490 (2023).

31. Linette, G. P. et al. Cardiovascular toxicity and titin cross-reactivity of affinity-enhanced T cells in myeloma and melanoma. Blood 122, 863–871 (2013).

32. Vázquez Torres, S. et al. De novo design of high-affinity binders of bioactive helical peptides. Nature 626, 435–442 (2024).

33. Baker, D. et al. De novo designed proteins neutralize lethal snake venom toxins. (2024).

34. Huber, F. et al. A comprehensive proteogenomic pipeline for neoantigen discovery to advance personalized cancer immunotherapy. Nat. Biotechnol. 1–13 (2024) doi:10.1038/s41587-024-02420-y.

35. Zhang, L. et al. Monoclonal antibody blocking the recognition of an insulin peptide-MHC complex modulates type 1 diabetes. Proc. Natl. Acad. Sci. U. S. A. 111, 2656–2661 (2014).

36. Johnson, M. et al. NCBI BLAST: a better web interface. Nucleic Acids Res. 36, W5–W9 (2008).

37. Reynisson, B., Alvarez, B., Paul, S., Peters, B. & Nielsen, M. NetMHCpan-4.1 and NetMHCIIpan-4.0: improved predictions of MHC antigen presentation by concurrent motif deconvolution and integration of MS MHC eluted ligand data. Nucleic Acids Res. 48, W449–W454 (2020).

38. Vilar, S., Cozza, G. & Moro, S. Medicinal Chemistry and the Molecular Operating Environment (MOE): Application of QSAR and Molecular Docking to Drug Discovery. Curr. Top. Med. Chem. 8, 1555–1572 (2008).

39. Labute, P. Protonate3D: assignment of ionization states and hydrogen coordinates to macromolecular structures. Proteins 75, 187–205 (2009).

40. Sabri Dashti, D., Meng, Y. & Roitberg, A. E. pH-Replica Exchange Molecular Dynamics in Proteins Using a Discrete Protonation Method. J. Phys. Chem. B 116, 8805–8811 (2012).

41. Case, D. A. et al. AmberTools. J. Chem. Inf. Model. 63, 6183–6191 (2023).

42. Fischer, A.-L. M. et al. The Role of Force Fields and Water Models in Protein Folding and Unfolding Dynamics. J. Chem. Theory Comput. 20, 2321–2333 (2024).

43. Jorgensen, W. L., Chandrasekhar, J., Madura, J. D., Impey, R. W. & Klein, M. L. Comparison of simple potential functions for simulating liquid water. J. Chem. Phys. 79, 926–935 (1983).

44. Gapsys, V. & de Groot, B. L. On the importance of statistics in molecular simulations for thermodynamics, kinetics and simulation box size. eLife 9, e57589 (2020).

45. Maier, J. A. et al. ff14SB: Improving the Accuracy of Protein Side Chain and Backbone Parameters from ff99SB. J. Chem. Theory Comput. 11, 3696–3713 (2015).

46. Doll, J. D., Myers, L. E. & Adelman, S. A. Generalized Langevin equation approach for atom/solid-surface scattering: Inelastic studies. J. Chem. Phys. 63, 4908–4914 (1975).

47. Adelman, S. A. & Doll, J. D. Generalized Langevin equation approach for atom/solid-surface scattering: General formulation for classical scattering off harmonic solids. J. Chem. Phys. 64, 2375–2388 (1976).

48. Åqvist, J., Wennerström, P., Nervall, M., Bjelic, S. & Brandsdal, B. O. Molecular dynamics simulations of water and biomolecules with a Monte Carlo constant pressure algorithm. Chem. Phys. Lett. 384, 288–294 (2004).

49. Roe, D. R. & Cheatham, T. E. I. PTRAJ and CPPTRAJ: Software for Processing and Analysis of Molecular Dynamics Trajectory Data. J. Chem. Theory Comput. 9, 3084–3095 (2013).

50. Harris, C. R. et al. Array programming with NumPy. Nature 585, 357–362 (2020).

51. Hunter, J. D. Matplotlib: A 2D Graphics Environment. Comput. Sci. Eng. 9, 90–95 (2007).

52. Saini, S. K. et al. Empty peptide-receptive MHC class I molecules for efficient detection of antigen-specific T cells. Sci. Immunol. 4, eaau9039 (2019).

53. Deichmann, M. et al. Engineered yeast cells simulating CD19+ cancers to control CAR T cell activation. 2023.10.25.563929 Preprint at 10.1101/2023.10.25.563929 (2024).

54. Mirdita, M. et al. ColabFold-Making protein folding accessible to all. (2021).

55. Fernández-Quintero, M. L. et al. Challenges in antibody structure prediction. mAbs (2023).

56. Sewell, A. K. Why must T cells be cross-reactive? Nat. Rev. Immunol. 12, 669–677 (2012).

57. Kabsch, W. XDS. Acta Crystallogr. D Biol. Crystallogr. 66, 125–132 (2010).

58. Winn, M. D. et al. Overview of the CCP 4 suite and current developments. Acta Crystallogr. D Biol. Crystallogr. 67, 235–242 (2011).

59. McCoy, A. J. et al. Phaser crystallographic software. J. Appl. Crystallogr. 40, 658–674 (2007).

60. Adams, P. D. et al. PHENIX : a comprehensive Python-based system for macromolecular structure solution. Acta Crystallogr. D Biol. Crystallogr. 66, 213–221 (2010).

61. Emsley, P. & Cowtan, K. Coot : model-building tools for molecular graphics. Acta Crystallogr. D Biol. Crystallogr. 60, 2126–2132 (2004).

62. Williams, C. J. et al. MolProbity: More and better reference data for improved all-atom structure validation. Protein Sci. 27, 293–315 (2018).

