## Supplementary Information for "*De novo* designed pMHC binders facilitate T cell induced killing of cancer cells"

### Extended Data

**Extended Data Table 1.** Common properties of pMHC binders surpassing the final filtering step in the design campaigns against the pMHC complex SLLMWITQC/HLA-A*02:01 (Melanoma shared cancer antigen).

| MHC | Peptide | No. of binders considered in final filter step | ipAE range [Å] | pLDDT_binder range [%] | Average ratio of predicted secondary structure conformation [%] | Average length [No. amino acids] |
|---|---|---|---|---|---|---|
| HLA-A*02:01 | SLLMWITQC | 44 | 4.5 - 6.9 | 92 - 97 | β sheets: 3.57%<br>Right-handed αhelices: 95.1 %<br>Left-handed α helices: 0.88 %<br>Rest: 0.45 % | 123 |

**Extended Data Table 2.** Common properties of pMHC binders surpassing the final filtering step in the design campaigns against the pMHC complex RVTDESILSY/HLA-A*01:01 (Melanoma neoantigen). The diversification of pMHC binder prior to in vitro screening as described in table 2 is not reflected in the data for this table.

| MHC | Peptide | No. of binders considered in final filter step | ipAE range [Å] | pLDDT_binder range [%] | Average ratio of predicted secondary structure conformation [%] | Average length [No. amino acids] |
|---|---|---|---|---|---|---|
| HLAA*01:01 | RVTDESILSY | 96 | 4.9 - 5.4 | 93.5 - 96.5 | β sheets: 3.03 %<br>Right-handed α helices: 95.3 %<br>Left-handed α helices: 1.5 %<br>Rest: 0.35 % | 124 |

### Supplementary Data

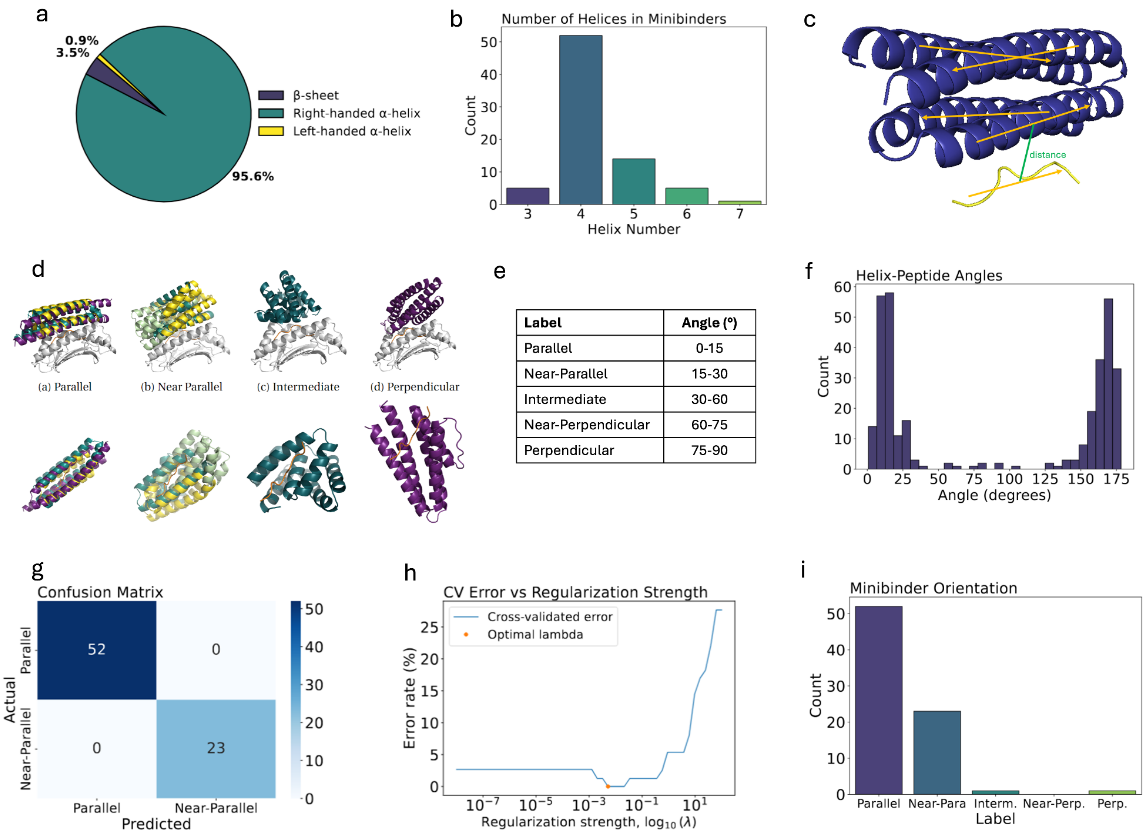


**Supplementary Figure S1. Structural and orientation analysis of miBds targeting the SLLMWITQC/HLA-A*02:01 NY-ESO-1 pMHC.** This covers analysis of 78 miBd designs with an ipAE below 7 Å and pLDDT above 90. **(a)** Secondary structure composition of miBds, where 95.6% of residues form right-handed α-helices, with minimal β-sheets (3.5%) and left-handed α-helices (0.9%). This highlights the dominance of right-handed α-helices in miBd designs. **(b)** Distribution of helices per miBd, with the majority containing four helices. This consistent feature contributes to the tightly packed arrangement of miBds. **(c)** Helix alignment relative to the peptide, illustrating the principal axes of helices and the peptide and the distance between. This alignment shows the encapsulation of the peptide by miBds helices. **(d)** 3D visualisations of miBds orientation labels relative to the peptide: parallel, near-parallel, intermediate, and perpendicular. **(e)** Angle ranges for each of the five orientations, from parallel (0–15°) to perpendicular (75–90°), was used to categorise miBds based on the average helix-peptide angle. **(f)** Distribution of all helix-peptide angles, illustrating an either parallel or anti-parallel favoured orientation, before adjusting for directional equivalency. **(g)** Confusion matrix showing the accuracy of the linear regression classifier for parallel and near-parallel orientations. The model achieved perfect accuracy, though its applicability and generalisability was limited due to the limited size and variation of the dataset. **(h)** Cross-validation error rates as a function of regularisation strength (log10 λ), identifying the optimal λ value. **(i)** Distribution of miBd orientation, with the majority classified as parallel or near-parallel. This shows a strong preferential miBd orientation towards the NY-ESO-1 peptide.

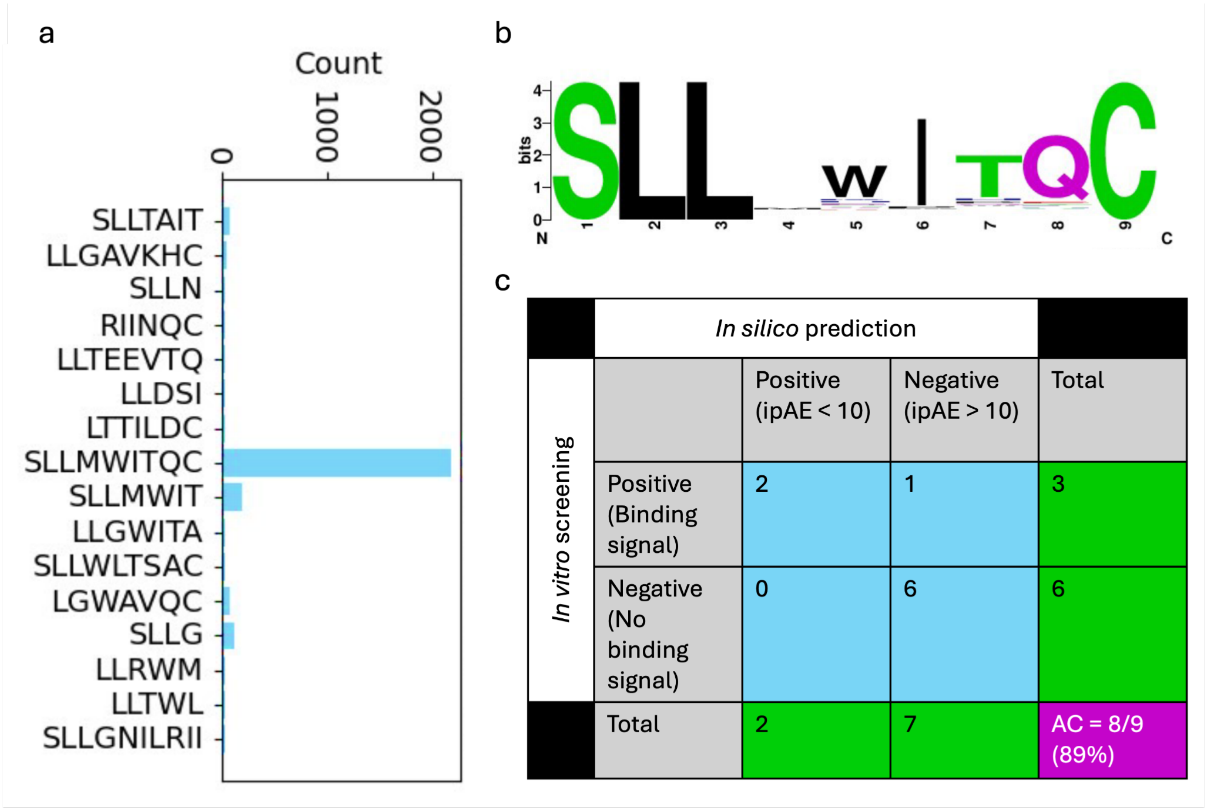


| | | *In silico* prediction | | |
|---|---|---|---|---|
| *In vitro* screening | | Positive (ipAE < 10) | Negative (ipAE > 10) | Total |
| | Positive (Binding signal) | 2 | 1 | 3 |
| | Negative (No binding signal) | 0 | 6 | 6 |
| | Total | 2 | 7 | AC = 8/9 (89%) |

**Supplementary Figure S2. Computational screening for potential cross-reactive binding of miBd against peptide variants of the SLLMWITQC NY-ESO-1 peptide presented on HLA-A*02:01. (a)** Total count of unique matching (identity >85%) sequences in the human proteome to 3610 single & double mutated peptides (of SLLMWITQC) utilising the web-based tool of NCBI pBLAST followed by filtering in NetMHCpan4.1 (accessible via DTU Health Tech Bioinformatics Services via https://services.healthtech.dtu.dk/services/NetMHCpan-4.1/). 'SLLGNILRI' obtained a ranking score within the highest 0.5-percentile and was considered as a peptide likely to be represented on the same HLA-A*02:01. **(b)** Sequence logo displaying sequence variation of the 1128 single and double mutated peptides (out of 3610) which obtained better or equal predicted affinity towards HLA-A*02:01 as SLLMWITQC. The residue in position 4, and to a lesser extent also the residues in positions 5, 7, and 8 were the least conserved residues with no or only a marginally dominating amino acid. **(c)** Comparison of in silico (Fig. 1c) and in vitro (Fig. 3 e-g) cross-reactivity screening for NY1-B04 on 9 peptide variants (SLLDFITQC, SLLDWIHQC, SLLDWIPQC, SLLDWITQC, SLLEHITQC, SLLEWITQC, SLLGNIRLI, SLLAWITQC, SLLMWITQV). The peptide variant SLLMYITQC was only screened in silico and, therefore, not considered alongside the original SLLMWITQC NY-ESO-1 peptide. The accuracy (AC) is defined as the sum of True negative and true positive predictions (assuming the in vitro results are the ground truth) divided by the total number of observations. The informative value is limited, due to the small data set size of only N = 9 and should only be regarded as a notable observation.

**Supplementary Table S3. List of gRNA, primers, pMB-LINK elements and peptides used in this study.** crRNA CD3ε was used for generating CD3 KO Jurkat cells, pMB-LINK is the CAR vector plasmid used for cloning all the miBd libraries and single constructs, primers were used for sequencing of the pMB_LINK, SLLMWITQC variant peptides were used for *in silico* and/or *in vitro* cross-panning.

| ELEMENT | Type | Sequence |
|---|---|---|
| crRNA CD3ε | DNA | GATGTCCACTATGACAATTG |
| pMB-LINK - CD8 leader sequence | DNA | ATGGCCCTCCCTGTCACCGCCCTGCTGCTTCCGCTGGCTCTTCT GCTCCACGCCGCTCGGCCC |
| pMB-LINK - BsmBI cloning sites | DNA | AGAGACGGTTGTGGAATTCTTCTACGTCTC |
| pMB-LINK - Linker | AA | GGGGSGGGGSGGGGS |
| pMB-LINK - CD8α hinge | AA | TTTPAPRPPTPAPTIASQPLSLRPEACRPAAGGAVHTRGLDFACDIYI WAPLAGTCGVLLLSLVITLYC |
| pMB-LINK - CD28 | AA | RSKRSRLLHSDYMNMTPRRPGPTRKHYQPYAPPRDFAAYRS |
| pMB-LINK - CD3ζ | AA | RVKFSRSADAPAYQQGQNQLYNELNLGRREEYDVLDKRRGRDPEM GGKPRRKNPQEGLYNELQKDKMAEAYSEIGMKGERRRGKGHDGLY QGLSTATKDTYDALHMQALPPR |
| pMB-LINK - GFP | AA | MVSKGEELFTGVVPILVELDGDVNGHKFSVSGEGEGDATYGKLTLKF ICTTGKLPVPWPTLVTTLTYGVQCFSRYPDHMKQHDFFKSAMPEGY VQERTIFFKDDGNYKTRAEVKFEGDTLVNRIELKGIDFKEDGNILGHK LEYNYNSHNVYIMADKQKNGIKVNFKIRHNIEDGSVQLADHYQQNTPI GDGPVLLPDNHYLSTQSALSKDPNEKRDHMVLLEFVTAAGITLGMDE LYK |
| pMB-LINK - cPPT-CTS sequence | DNA | TTTTAAAAGAAAAGGGGGGATTGGGGGGTACAGTGCAGGGGAAA GAATAGTAGACATAATAGCAACAGACATACAAACTAAAGAATTACA AAAACAAATTACAAAAATTCAAAATTTT |
| pMB-LINK - EF-1α promoter | DNA | GGCTCCGGTGCCCGTCAGTGGGCAGAGCGCACATCGCCCACAG TCCCCGAGAAGTTGGGGGGAGGGGTCGGCAATTGAACCGGTGC CTAGAGAAGGTGGCGCGGGGTAAACTGGGAAAGTGATGTCGTGT ACTGGCTCCGCCTTTTTCCCGAGGGTGGGGGAGAACCGTATATA AGTGCAGTAGTCGCCGTGAACGTTCTTTTTCGCAACGGGTTTGCC GCCAGAACACAGGTAAGTGCCGTGTGTGGTTCCCGCGGGCCTG GCCTCTTTACGGGTTATGGCCCTTGCGTGCCTTGAATTACTTCCA CCTGGCTGCAGTACGTGATTCTTGATCCCGAGCTTCGGGTTGGA AGTGGGTGGGAGAGTTCGAGGCCTTGCGCTTAAGGAGCCCCTTC GCCTCGTGCTTGAGTTGAGGCCTGGCCTGGGCGCTGGGGCCGC CGCGTGCGAATCTGGTGGCACCTTCGCGCCTGTCTCGCTGCTTT CGATAAGTCTCTAGCCATTTAAAATTTTTGATGACCTGCTGCGACG CTTTTTTTCTGGCAAGATAGTCTTGTAAATGCGGGCCAAGATCTG CACACTGGTATTTCGGTTTTTGGGGCCGCGGGCGGCGACGGGG CCCGTGCGTCCCAGCGCACATGTTCGGCGAGGCGGGGCCTGCG |

| | | |
|---|---|---|
| | | AGCGCGGCCACCGAGAATCGGACGGGGGTAGTCTCAAGCTGGC CGGCCTGCTCTGGTGCCTGGCCTCGCGCCGCCGTGTATCGCCC CGCCCTGGGCGGCAAGGCTGGCCCGGTCGGCACCAGTTGCGTG AGCGGAAAGATGGCCGCTTCCCGGCCCTGCTGCAGGGAGCTCA AAATGGAGGACGCGGCGCTCGGGAGAGCGGGCGGGTGAGTCAC CCACACAAAGGAAAAGGGCCTTTCCGTCCTCAGCCGTCGCTTCAT GTGACTCCACTGAGTACCGGGCGCCGTCCAGGCACCTCGATTAG TTCTCGTGCTTTTGGAGTACGTCGTCTTTAGGTTGGGGGGAGGG GTTTTATGCGATGGAGTTTCCCCACACTGAGTGGGTGGAGACTGA AGTTAGGCCAGCTTGGCACTTGATGTAATTCTCCTTGGAATTTGC CCTTTTTGAGTTTGGATCTTGGTTCATTCTCAAGCCTCAGACAGTG GTTCAAAGTTTTTTTCTTCCATTTCAGGTGTCGTGA |
| pMB-LINK - Woodchuck Hepatitis Virus (WHV) Post-transcriptional Regulatory Element (WPRE) | DNA | AATCAACCTCTGGATTACAAAATTTGTGAAAGATTGACTGGTATTC TTAACTATGTTGCTCCTTTTACGCTATGTGGATACGCTGCTTTAAT GCCTTTGTATCATGCTATTGCTTCCCGTATGGCTTTCATTTTCTCC TCCTTGTATAAATCCTGGTTGCTGTCTCTTTATGAGGAGTTGTGGC CCGTTGTCAGGCAACGTGGCGTGGTGTGCACTGTGTTTGCTGAC GCAACCCCCACTGGTTGGGGCATTGCCACCACCTGTCAGCTCCT TTCCGGGACTTTCGCTTTCCCCCTCCCTATTGCCACGGCGGAACT CATCGCCGCCTGCCTTGCCCGCTGCTGGACAGGGGCTCGGCTG TTGGGCACTGACAATTCCGTGGTGTTGTCGGGGAAGCTGACGTC CTTTCCTTGGCTGCTCGCCTGTGTTGCCACCTGGATTCTGCGCG GGACGTCCTTCTGCTACGTCCCTTCGGCCCTCAATCCAGCGGAC CTTCCTTCCCGCGGCCTGCTGCCGGCTCTGCGGCCTCTTCCGCG TCTTCGCCTTCGCCCTCAGACGAGTCGGATCTCCCTTTGGGCCG CCTCCCCGC |
| MB_GGinterm_fwd_2 | DNA | TTCATTCTCAAGCCTCAGACAGT |
| MB_GGinterm_rev_2 | DNA | GATATCGCAGGCGAAGTCAAGA |
| Mutated SLLMWITQC peptide (for *in silico* and *in vitro* cross-panning) | AA | SLLDFITQC |
| Mutated SLLMWITQC peptide (for *in silico* and *in vitro* cross-panning) | AA | SLLDWIHQC |
| Mutated SLLMWITQC peptide (for *in silico* and *in vitro* cross-panning) | AA | SLLDWIPQC |
| Mutated SLLMWITQC peptide (for *in silico* and *in vitro* cross-panning) | AA | SLLDWITQC |

| | | |
|---|---|---|
| Mutated SLLMWITQC peptide (for *in silico* and *in vitro* cross-panning) | AA | SLLEHITQC |
| Mutated SLLMWITQC peptide (for *in silico* and *in vitro* cross-panning) | AA | SLLEWITQC |
| Mutated SLLMWITQC peptide (for *in silico* and *in vitro* cross-panning) | AA | SLLGNIRLI |
| Mutated SLLMWITQC peptide (for *in silico* cross-panning) | AA | SLLMYITQC |
| Mutated SLLMWITQC peptide (for *in silico* and *in vitro* cross-panning) | AA | SLLAWITQC |
| Mutated SLLMWITQC peptide (for *in silico* and *in vitro* cross-panning) | AA | SLLMWITQV |
